# Variation in intraspecific demography drives localised concordance but species-wide discordance in responses to Plio-Pleistocene climatic change

**DOI:** 10.1101/2021.09.21.461180

**Authors:** Sean James Buckley, Chris Brauer, Peter J. Unmack, Michael P. Hammer, Luciano B. Beheregaray

## Abstract

Understanding how species biology may facilitate resilience to climate change remains a critical factor in detecting and protecting species at risk of extinction. Many studies have focused on the role of particular ecological traits in driving species responses, but less so on demographic history and levels of standing genetic variation. We used environmental and genomic datasets to reconstruct the phylogeographic histories of two ecologically similar and largely co-distributed freshwater fishes to assess the degree of concordance in their responses to Plio-Pleistocene climatic changes. Although several co-occurring populations demonstrated concordant demographic histories, idiosyncratic population size changes were found at the range edges of the more spatially restricted species. Discordant responses between species were associated with low standing genetic variation in peripheral populations. This might have hindered adaptive potential, as documented in recent population declines and extinctions of the two species. Our results highlight both the role of spatial scale in the degree of concordance in species responses to climate change, and the importance of standing genetic variation in facilitating range shifts. Even when ecological traits are similar between species, long-term genetic diversity and historical population demography may lead to discordant responses to ongoing and future climate change.

## Introduction

Understanding how or whether species may be able to adapt to current and future climatic changes is critical for conservation management of threatened taxa [1]. However, predicting the susceptibility and extent of species loss due to climate change remains a challenge. To this end, many studies have instead sought to determine ecological traits that may confer resilience or susceptibility to climate change across various taxa [2]. Ecological and physiological traits such as thermal tolerance and dispersal capacity have been shown to be critical in driving adaptation to climatic changes [3, 4]. Demographic and genetic traits such as population size, stability and standing genetic variation (SGV) are however also important in facilitating adaptation to new environmental stressors [5], and likely play a major role in species responses to climate change [6-8].

From a genetic perspective, adaptation to novel climatic conditions more often relies upon SGV than *de novo* mutations [9-11]. The degree to which SGV is maintained within species or populations varies substantially across taxa and is influenced by a combination of demographic, ecological and environmental factors. For example, populations occurring at the edge of a species range often have lower connectivity and genetic diversity than their more central counterparts [12], including reduced diversity in climate-associated genes [13]. In marginal populations, persistence is driven by the balance of the steepness of the selective environment and the effectiveness of selection relative to genetic drift [14]. These components may contrast with the core of the distribution, where larger carrying capacities and SGV allow populations to persist closer to their selective optimum [15]. Thus, the interaction and spatial variability of neutral (demographic) and adaptive (ecological) traits are critically important in understanding how species ranges may shift under climate change [16].

Understanding factors underlying species responses to historical climatic fluctuations provides an empirical framework for determining how species may respond to current and future environmental changes [17]. Extending phylogeographic analyses from taxon-specific studies to assessments of how species assemblages have responded to past climatic changes provides an approach to estimating the ubiquity of species responses [18]. Similar species responses (concordance) across disparate taxa often indicate that shared ecological traits underlie the response [19], or demonstrate the ubiquity in impact of the environmental change in question [20]. Contrastingly, idiosyncratic responses (discordance) are often attributed to variation in species-specific ecological traits [21]. However, intraspecific variation in demography may lead to spatial variation in the degree of concordance, even across ecologically similar species. For example, the interactive role of demography and adaptive potential may lead to intraspecific variation at local scales, even if species-wide patterns are concordant across taxa or vice versa [22, 23]. These patterns may be reflected within species range shifts over time, where intraspecific variation in demographic or ecological traits at range margins may drive interspecific discordance in species responses to environmental change.

Biogeographic regions that experienced major environmental change in the past are particularly useful for studying species responses to climate change. In this regard, the southeast Australian temperate zone provides a model region to test how species have responded to major environmental changes such as aridification and eustatic changes. Mainland Australia has experienced significant environmental changes since the late Miocene, which heralded the onset of major aridification [24]. Other than a brief humid period during the Pliocene [25], this aridification intensified into the Pleistocene. While glacial periods in this region were not directly associated with the formation of glaciers, major changes in precipitation and temperature shifted ecosystems towards more arid conditions [26]. Concordantly, glacial maxima also drove eustatic changes, expanding much of the continental shelf as sea levels dropped [27]. The complex environmental history in southeast Australia, and its role on the evolution of temperate species, has been demonstrated by a number of phylogeographic studies (e.g. [28, 29]).

Freshwater-dependent species are important indicators of historical environmental changes given their reliance on suitable habitat and often limited capacity for dispersal [30]. Within temperate southeast Australia, the often co-distributed southern (*Nannoperca australis*) and Yarra (*N. obscura*) pygmy perches provide an ideal comparative study system. Both species possess highly similar morphology, reproductive biology, salinity tolerance and habitat preferences, and also display similar patterns of metapopulation structure [31-35]. Both species have low dispersal capacity with little to nil contemporary connectivity among catchments [33, 35]. Both species are relatively old (e.g. their lineages diverged around 13 million years ago [36]) and show strong population structure, with two evolutionarily significant units (ESUs) separating coastal and inland (Murray-Darling Basin) populations in *N. australis* [28], and two clades each containing two ESUs in *N. obscura* [37]. Given their isolated populations, it is expected that their long-term persistence along landscapes depends on spatial variation of locally adaptive traits. This hypothesis is consistent with studies of *N. australis* that show that patterns of adaptation in traits related to reproductive fitness [38, 39], in levels of adaptive genetic diversity [34] and in variance of gene expression [40] are strongly associated with hydroclimatic gradients.

Despite their ecological similarities, the two species demonstrate marked differences in conservation status, genetic diversity and total distribution range. While both species are of conservation concern (*N. australis* as Near Threatened and *N. obscura* as Endangered) within the IUCN Red List [41] and in state conservation legislation, *N. obscura* is considered at higher risk due to their narrow range and extremely low genetic diversity [35, 42]. These factors are implicated in the local extirpation of *N. obscura* within the Murray-Darling Basin in the last five years, following failed reintroductions after a large-scale drought impacted the region [41]. The relatively low genetic diversity of *N. obscura* is not thought to be the result of any particularly severe past bottleneck [42], complicating determining factors underlying this disparity. Additionally, it remains unclear whether the historical absence of *N. obscura* in some regions where *N. australis* is found is the result of historical local extinctions or a failure to initially colonise the habitat.

Here, we applied a comparative phylogeographic framework to explore the relative roles of ecological and demographic traits on evolutionary history. We used genomic datasets to estimate genetic diversity, phylogenetic relationships and demographic history of these two freshwater fishes, in conjunction with species distribution modelling. Then, we statistically evaluated regional concordance across co-occurring populations to assess whether the species shared demographic responses to Pleistocene glacial cycles. We predicted that evolutionary patterns, demographic histories and distribution changes would be concordant across the two species if ecological factors played a relatively strong role in determining species responses to past climatic changes, with current differences owing to more recent factors.

Contrastingly, discordant histories would indicate that genetic diversity and demography played a relatively larger role and underpinned their contemporary differences in conservation status. Our framework also includes differentiation of local-scale (population-level) and broad-scale (species-level) responses to assess the role of intraspecific patterns in driving lineage responses.

## Methods

### Sample collection and genomic library preparation

The distribution of both species spans the southwest Victoria biogeographic province and the lower reaches of the Murray-Darling Basin [31]. *Nannoperca australis* is more widely distributed and is also found across eastern Victorian drainages, northern Tasmania and the upper reaches of the southern Murray-Darling Basin [43]. The final sample contains all known genetically distinct populations (including recently extirpated populations) across their full co-distributed range (electronic supplementary material, Table S1). This equals to seven populations of *N. obscura* and nine populations of *N. australis* occurring across all major drainages of the region (Figure 1). An additional 10 and 15 *N. obscura* and *N. australis* (respectively) from Lake Alexandrina within the lower Murray-Darling Basin were also included for more targeted demographic reconstruction of these populations. For phylogenetic analyses, five samples of a sister species (*Nannoperca vittata*) were included as outgroup [36]. Specimens were collected using electrofishing, dip-, fyke- or seine-netting. Either the caudal fin or the entire specimen was stored at −80°C at the South Australian Museum, or in 99% ethanol at Flinders University.

**Figure 1:**
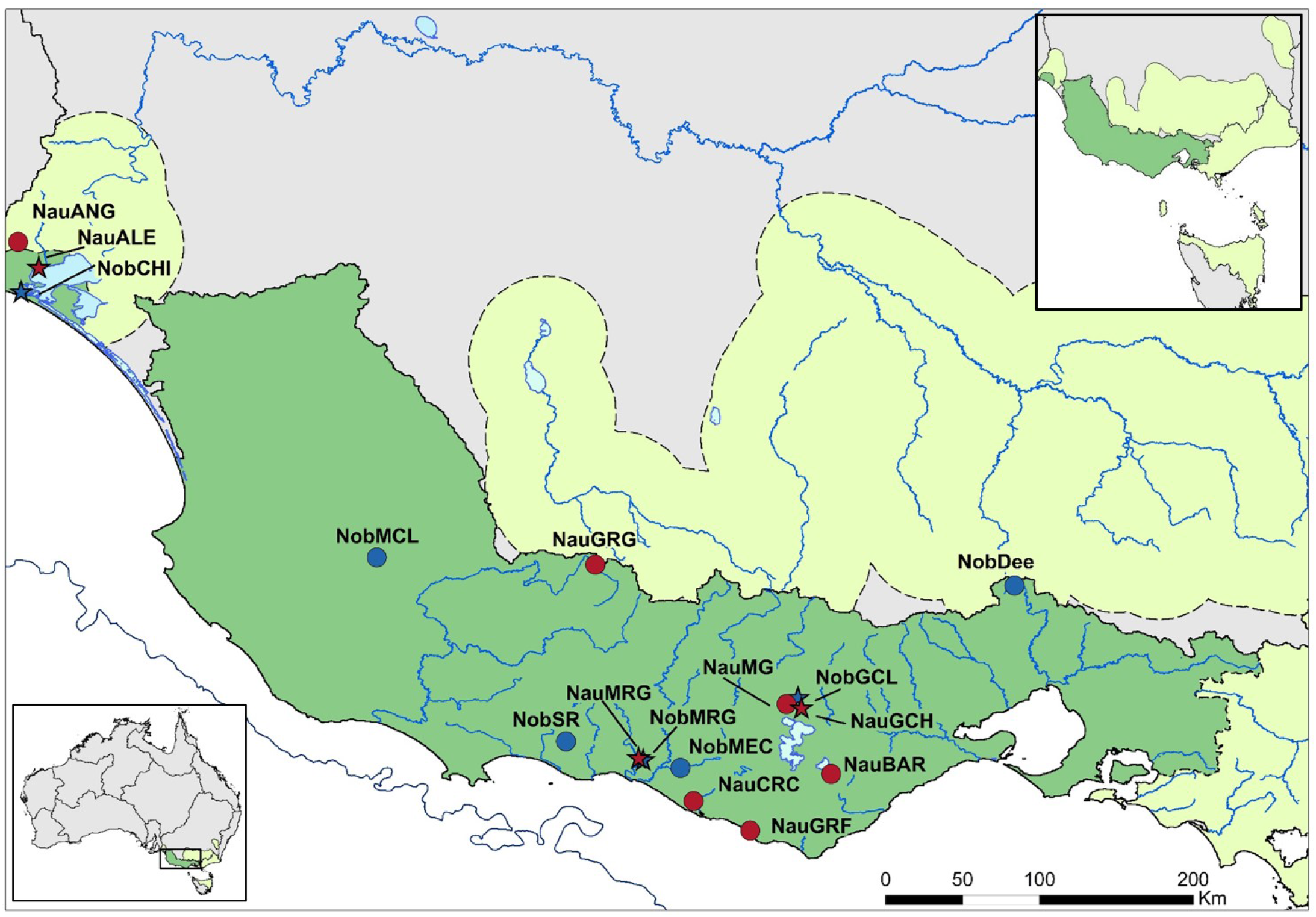
Contemporary distribution and sampling map for *N. australis* and *N. obscura. Nannoperca australis* sampling sites are indicated in red, and *N. obscura* sites in blue. The distribution of *N. australis* is indicated with light green shading and dashed borders, with the distribution of *N. obscura* (also the region of co-occurrence) in darker green. The solid black line indicates the boundary of major drainage basins, and the dark blue line demonstrates the approximate shoreline during glacial maxima. Bottom left inset depicts study region and major drainage basins in Australia. Top right inset depicts the full extent of species distributions.

DNA was extracted from muscle tissue or fin clips using a modified salting-out method [44] or a Qiagen DNeasy kit (Qiagen Inc., Valencia, CA, USA). Genomic DNA was checked for quality using a spectrophotometer (NanoDrop, Thermo Scientific), integrity using 2% agarose gels, and quantity using a fluorometer (Qubit, Life Technologies). The ddRAD (double digest restriction-site associated DNA) genomic libraries were prepared in-house at the Molecular Ecology Lab of Flinders University following [34]. The majority of the samples (56/98) were paired-end sequenced on an Illumina HiSeq 2000 at Genome Quebec (Montreal, Canada). The remaining samples were single-end sequenced on an Illumina HiSeq 2500 at the South Australia Health and Medical Research Institute (SAHMRI).

### Sequence filtering, alignment and SNP calling

Sequences were demultiplexed using the ‘process_radtags’ module of Stacks 1.29 [45], allowing up to 2 mismatches in barcodes. Barcodes were removed and sequences trimmed to 80 bp to remove low-quality bases from the end of the reads. Trimmed reads were aligned using PyRAD 3.0.6 [46], and further cleaned by removing reads with >5 bp with a Phred score < 20. Loci were retained if they occurred in at least ∼80% of samples (22 in *N. obscura*; 30 in *N. australis)* within the phylogenetic datasets.

For some population-specific analyses (e.g. some genetic diversity measures and coalescent-based demographic histories), we subsampled our ddRADseq data with loci re-aligned and SNPs called separately for each population (excluding those with *n* < 3 due to low sample size) using PyRAD. This was done as most SNPs called at the species level were monomorphic within individual populations and would have significant effects on downstream analyses [47]. Only loci present in all individuals were kept to prevent missing data from biasing the site-frequency spectrum (SFS), used in all demographic analyses [48].

### Contemporary genetic diversity

Summaries of population-level genetic diversity parameters (allelic richness and gene diversity) were compared across populations within species, and between species, using the R package *hierfstat* [49]. Given uneven sample sizes, rarefaction was used (*n* = 4) to estimate mean values per locus per population. Due to the larger sample sizes available for Lake Alexandrina populations, genetic diversity parameters were also calculated using *n* = 15 rarefaction and loci aligned separately within each Lake Alexandrina population. For these populations, we also calculated effective population size (*Ne*) using NeEstimator [50] and a minor allele frequency threshold of 0.02. Additionally, nucleotide diversity (π) within each population was estimated using dnaSP 6.1 [51]. Differences in population means of genetic diversity parameters between the two species were statistically evaluated using t-tests (two-tailed t-test or Wilcox test).

### Phylogenetic and historical migration analyses

Maximum likelihood (ML) phylogenies of each species were estimated using RAxML 8.2.11 [52] with the concatenated ddRAD alignments to estimate evolutionary relationships. Phylogenies were estimated under the GTR-GAMMA model of evolution and 1,000 RELL bootstraps for each species. Additionally, we estimated gene trees for each RAD locus using IQ-TREE2 [53] to account for genome-wide heterogeneity and incomplete lineage sorting. Gene and site concordance factors [54] were estimated by comparing individual gene trees to the concatenated RAxML tree.

As historical migration may impact the topology of a phylogenetic tree, we also used TreeMix [55] to infer historical population connectivity. We iteratively increased the number of migrations from 0 – *n* for each species (nine in *N. australis;* seven in *N. obscura*) and evaluated the fit of each tree based on the standard error of the covariance matrix. We further assessed the fit of the trees by calculating the percentage of variance explained per model (https://github.com/wlz0726/Population_Genomics_Scripts/tree/master/03.treemix). The best supported number of migrations was determined by the asymptote of the likelihood, where additional migrations did not substantially increase model likelihood.

### Comparative demographic inference

Long-term demographic histories for all populations were estimated using stairway plots and the SFS. One-dimensional SFS were calculated for each independent population alignment using an in-house script. Stairway plots were estimated assuming a mutation rate of 10^−8^ mutations per site per generation, and a generation time of one year for both species [42, 56]. Although both species reproduce annually, most individuals do not live beyond one to two years in the wild [56]. We then analysed co-distributed populations of *N. australis* and *N. obscura* under two coalescent frameworks to statistically evaluate the degree of concordant demographic history. The populations of Gnarkeet Creek (NauGCH and NobGCL), Merri River (NauMRG and NobMRG) and Lake Alexandrina (NauALE and NobCHI) were selected based on their contemporary co-occurrence and to represent the geographic range of the overlap in species distributions (Figure 1).

We first used FastSimCoal2 [57] to simulate model-based demographic histories over the last 30 Kyr. Simulations were conducted for each population under five different single population demographic scenarios (electronic supplementary material, Figure S1). Parameters were estimated using 40 optimisation cycles with 500,000 simulations per scenario, with the fit of the models estimated using Akaike Information Criterion (AIC) and Akaike weights (electronic supplementary material, Methods and Table S2). Confidence intervals for the parameters specified in the best supported demographic model per population were estimated by simulating 100 SFS and re-simulating point estimates using 500,000 iterations per SFS.

Additionally, we ran co-demographic model-based simulations using the aggregate site frequency spectrum (aSFS) and hierarchical approximate Bayesian computation in Multi-DICE [58] to determine if demographic histories were congruent across co-distributed populations. A single model of exponential growth followed by exponential decline was applied to all populations using broad uniform priors (electronic supplementary material, Methods and Figure S2), based on results from FastSimCoal2 (see xResults). We first tested the proportion of co-contracting taxa (ξ), and then fixed this hyperparameter to better explore the remaining parameters. A “leave-one-out” approach using 50 pseudo-observed datasets was used to generate a confusion matrix, with the most likely proportion of co-contracting taxa determined using the top 1,500 simulations and Bayes Factors. Parameters were estimated using 1.5 million coalescent simulations and posterior distributions estimated using the top 100 simulations and the *abc* R package [59]. We further tested whether demographic syndromes were broadly consistent across populations by comparing the posterior distributions for *Ne* and bottleneck strength (ε).

### Contemporary and paleoclimatic environmental modelling

Species distribution models (SDMs) were estimated using an ensemble modelling approach within biomod2 [60]. We estimated SDMs for both species across eleven time slices ranging from contemporary conditions to the Pliocene using the PaleoClim database [61]. Occurrence records for both species were obtained from a combination of sampled sites within this and past studies [28, 35, 36], as well as from the Atlas of Living Australia (http://www.ala.org.au/). We filtered the occurrence data to reduce the impact of spatial autocorrelation, resulting in final datasets of 1,021 and 163 observations for *N. australis* and *N. obscura*, respectively (electronic supplementary material, Methods).

We selected eight non-correlated environmental variables for estimating species distributions (electronic supplementary material, Table S3). These were annual mean temperature (Bio1), mean diurnal range (Bio2), isothermality (Bio3), temperature seasonality (Bio6), mean temperature of the wettest quarter (Bio8), mean temperature of the driest quarter (Bio9), annual precipitation (Bio12) and precipitation seasonality (Bio15). For the three oldest time periods, Bio2, Bio3 and Bio6 were unavailable and thus not included within the projections. SDMs were estimated using MaxEnt, random forest and generalised linear models, and an ensemble model generated per time period using the weighted mean of all models. All models were evaluated using both the relative operating characteristic and the true skill statistic. We quantitatively assessed the relative stability of species distributions over time by estimating the mean and standard deviation of suitability over time for each species. Differences in distributional ranges between species across time were estimated by converting SDMs to binary presence-absence maps based on the minimum suitability of the top 90% of putative occurrences per model (electronic supplementary materials, Methods).

## Results

### Bioinformatics

We obtained 21,051 ddRAD loci containing 53,334 filtered SNPs for *N. obscura* and 19,428 ddRAD loci containing 69,264 filtered SNPs for *N. australis*, with low missing data in both alignments (electronic supplementary material, Figure S1). Genetic diversity differed remarkably between the two species, with allelic richness, gene diversity, nucleotide diversity and number of SNPs per population alignment being significantly higher (p ≤ 0.01) in *N. australis* (electronic supplementary material, Table S1).

### Phylogenetic analysis

Phylogenetic analysis of both datasets returned a highly supported phylogenetic tree for each species. Site concordance factors broadly supported these patterns, although gene concordance factors were low across both trees (electronic supplementary material, Figures S4–S6) – this is not unexpected when gene trees are estimated from short and relatively uninformative individual loci [54]. For southern pygmy perch, the topology of this phylogenetic tree mirrored the geographic range of the samples, with a clear division between the Murray-Darling Basin ESU and the coastal ESU within the tree (Figure 2A). Within the coastal clade, populations diverged in a longitudinal manner, with eastern populations as the most recently diverged. In contrast, the phylogenetic tree for *N. obscura* did not demonstrate the same precise patterns, with populations not diverging in an exactly longitudinal manner. However, this was driven by a single outlier population (NobMEC).

**Figure 2:**
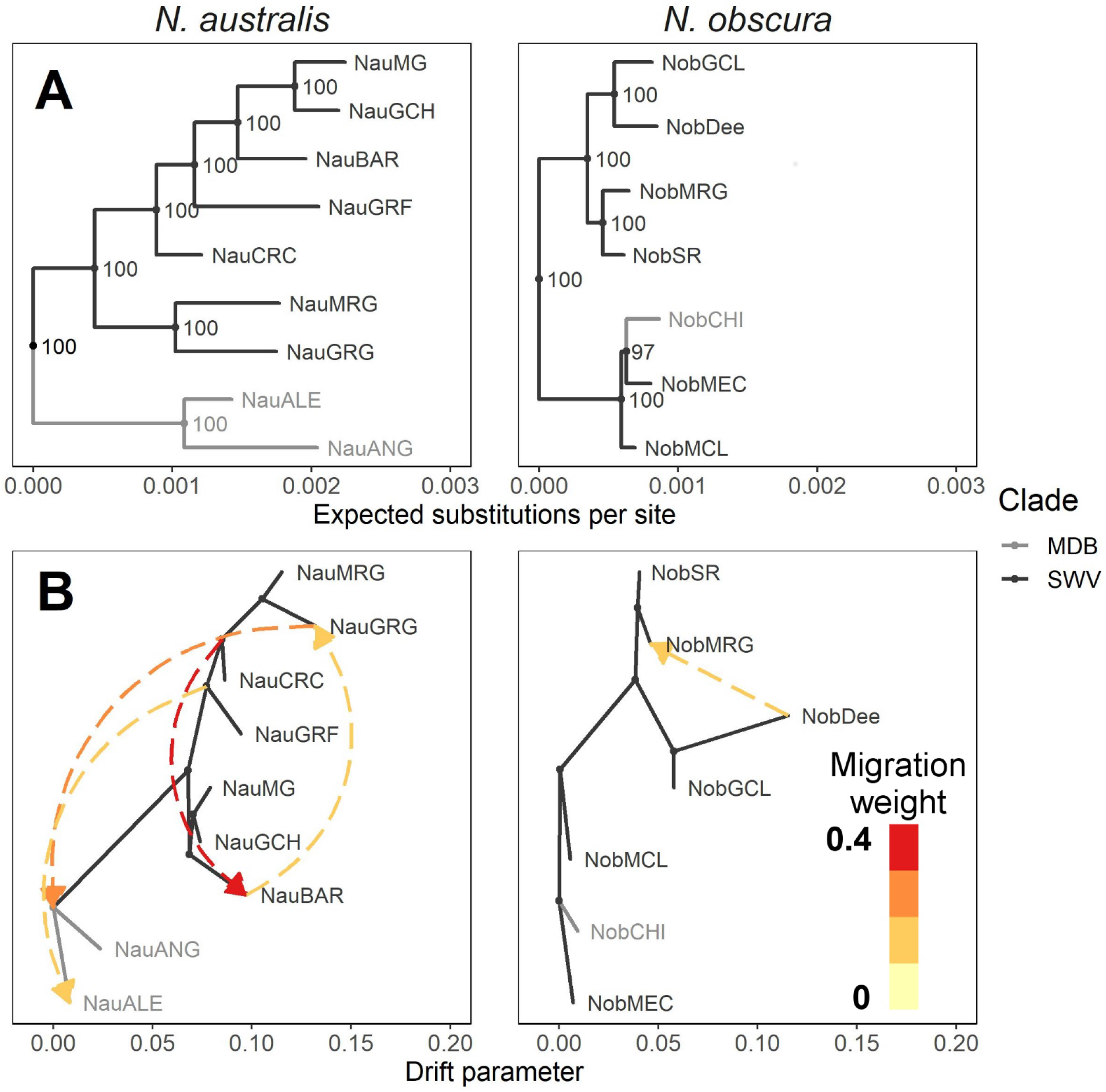
Phylogenetic histories and migration patterns in *N. australis* and *N. obscura*. **A:** Maximum likelihood phylogenetic trees based on ddRAD loci. Populations were reciprocally monophyletic and so were collapsed to the population level for simplicity. Both trees were rooted using *N. vittata* as the outgroup, which was dropped for visualisation. Node values show bootstrap support. Branch colours indicate the drainage basin of origin for each population or clade. **B:** Best supported ancestral migration patterns inferred using TreeMix based on SNP datasets. All displayed migrations were statistically significant (p < 0.05). Arrows denote the direction of inferred migrations, with the colour indicating their relative weights.

TreeMix inferred a greater number of migration events within *N. australis* (four) than *N. obscura* (one event) (Figure 2B; electronic supplementary material, Figure S7). Within *N. australis*, migrations were inferred both across populations of the coastal lineage as well as into the Murray-Darling Basin. The strongest migrations were between eastern coastal populations, and from the ancestor of the westernmost coastal population into the ancestor of the Murray-Darling Basin lineages. For *N. obscura*, the single migration inferred suggested historical gene flow from the easternmost population to a more central population. Trees and migration edges for both species were well supported by covariance matrices, with low pairwise residuals (electronic supplementary material, Figure S8) and standard errors of < 1 for any given population for both species (electronic supplementary material, Figure S9).

### Comparative demography

Stairway plots demonstrated broadly similar demographic histories across the two species, with most populations relatively stable or declining slightly over the last 1 Mya (Figure 3A). Populations within both species demonstrated variable demographic histories, although populations of *N. australis* appeared generally more stable over time. Both Lake Alexandrina populations (NobCHI and NauALE) showed significant historical increases in *Ne* >200 Kya, and long-term stable population sizes following this expansion.

**Figure 3:**
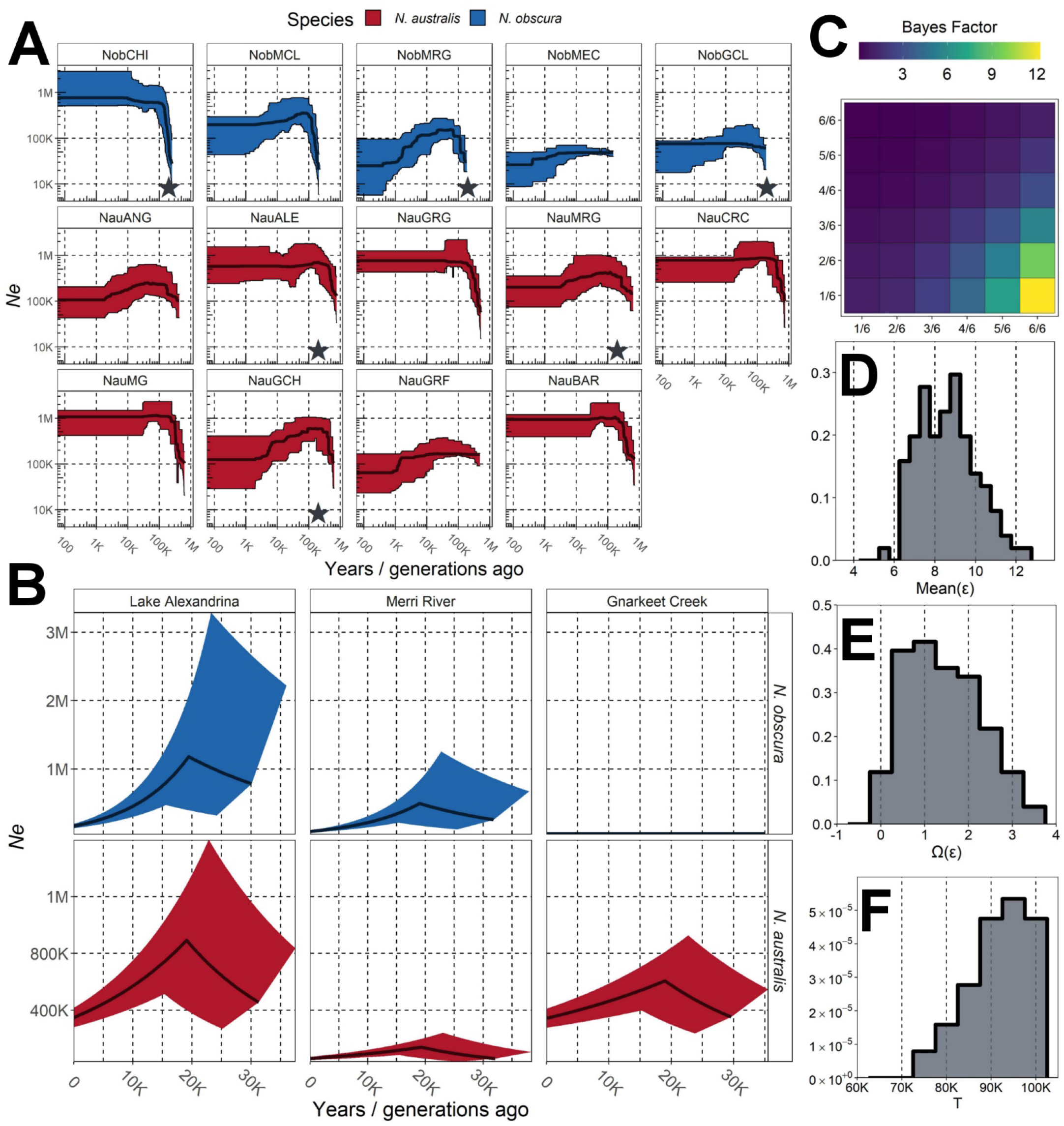
Demographic histories of *N. australis* and *N. obscura* populations. **A:** Stairway plot reconstructions of demographic history. Inset stars indicate co-occurring populations which were further explored within a codemographic framework. Populations are arranged from westernmost to easternmost within each species. **B:** Most likely individual demographic histories for co-occurring *N. australis* and *N. obscura* populations over the Pleistocene, simulated using FastSimCoal2. Thick dark lines indicate mean *Ne* over time, calculated based on the means of current *Ne*, rates of change and timing of switching rates (see Supplementary Material). Shaded areas indicate 95% confidence intervals based on the 97.5% and 2.5% probability estimates for the same parameters. **C:** Bayes Factor matrix of the proportion of populations showing synchronised bottlenecks (ξ) within a co-demographic model using Multi-DICE. Each cell compares the model in the column with the model in the row, with brighter colours indicating greater support for the column. **D:** Posterior distribution of mean bottleneck strength (ε) across all six populations. **E:** Posterior distribution of dispersion index of bottleneck strength (Var(ε)/Mean(ε)) across all six populations. **F:** Posterior distribution of the timing of the bottleneck event, in generations/years.

Most populations chosen for comparative analysis demonstrated fluctuating demographic histories (Figure 3B), with a period of pre-LGM (Last Glacial Maximum) growth followed by a post-LGM decline (electronic supplementary material, Figure S10). Only the eastern *N. obscura* population (NobGCL) contrasted this pattern, with a model of low but constant population size more supported than other demographic histories (Model 3). Strong post-glacial declines were present in Lake Alexandrina populations of both species, with weaker declines in the more eastern population pairs.

A confusion matrix suggested that the co-demographic model was more likely to infer fully synchronous (ξ = 1) or fully asynchronous (ξ = 0.167) co-contractions over intermediate proportions of taxa (electronic supplementary material, Figure S11). Despite this, Bayes Factors supported a fully synchronous model over more asynchronous models, and so ξ was fixed to 1 to better explore other parameters (Figure 3C). Contemporary population sizes were inferred to be relatively small across all populations with relatively weak post-glacial bottleneck strength (Figure 3D; electronic supplementary material, Table S5). These bottlenecks were similar in magnitude across populations, as indicated by low values of the dispersion index (Figure 3E). However, Multi-DICE did not recover the same timing of the bottleneck, possibly due to relatively low resolution within the aSFS (Figure 3F). Overall, these results support a widespread and concordant bottleneck across the six co-distributed populations.

### Species distribution modelling

Comparing the SDMs of the two species indicated much greater maximum distribution and variation in distributional range in *N. australis* than in *N. obscura. Nannoperca obscura* demonstrated long-term isolation to a relatively small region of southwest Victoria, whilst *N. australis* demonstrated a significant range expansion event throughout the early Pleistocene with a more recent contraction in the Holocene (electronic supplementary material, Figure S12). Despite these differences, both species maintained a shared climatic refugium in southwest Victoria, highlighted by a region of high mean suitability in both species (Figure 4B).

**Figure 4:**
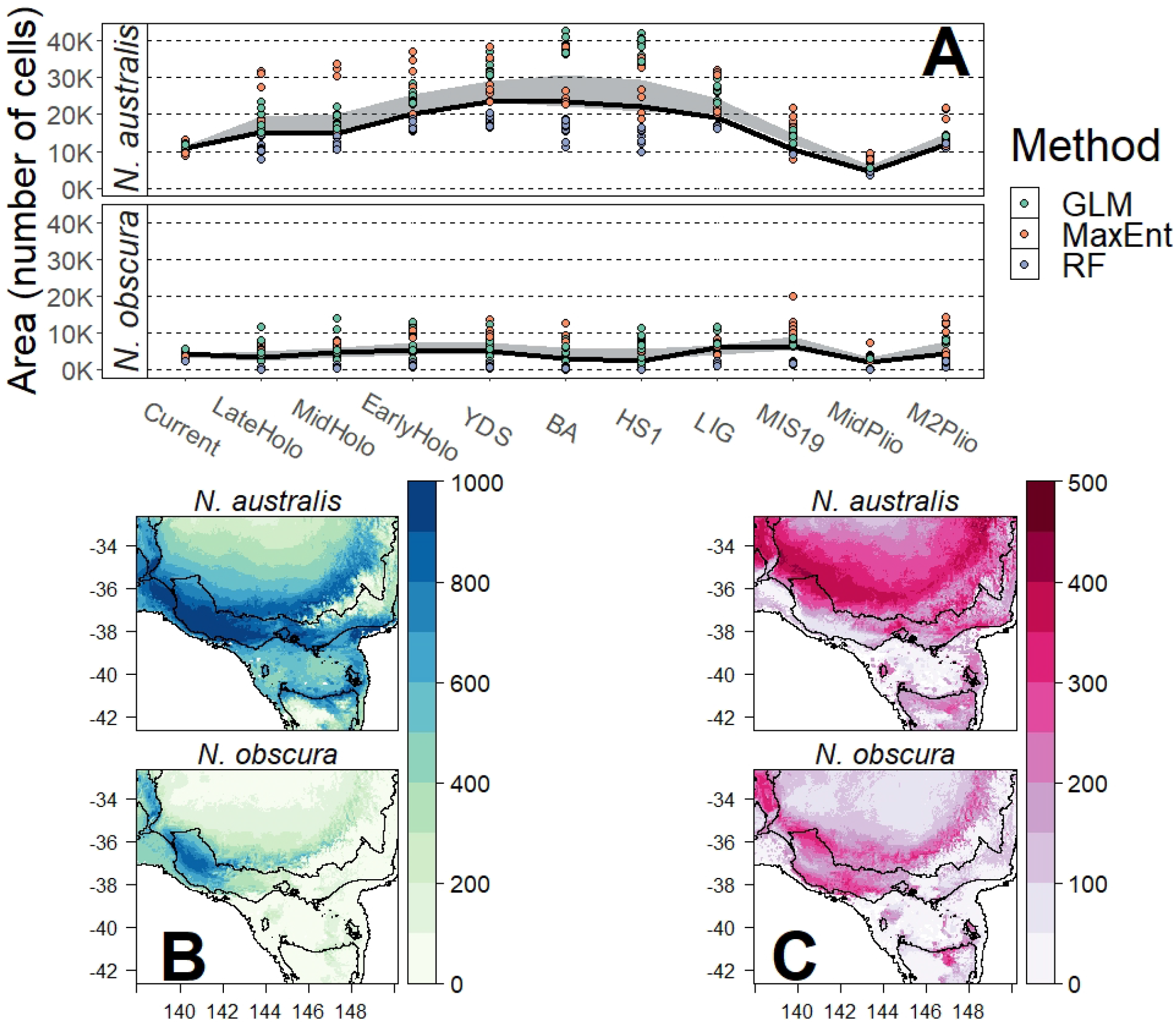
Comparisons of summaries of distributional changes over eleven time periods spanning the Plio-Pleistocene. **A:** Distribution extent per species. Individual models are indicated by points, with SDM method indicated by colour. The 95% confidence interval across all individual models is shown by the pale blue ribbon. The ensemble model is represented by a solid black line. **B:** Mean cell suitability across all time periods. **C:** Variation (standard deviation) in cell suitability across all time periods.

Comparisons across the different methods indicated that RandomForest was more conservative in estimating area (Figure 4A). While there was significant variation in estimated area across the different methods, ensemble models approximately captured the mean of all models. *Nannoperca australis* demonstrated significantly larger distributions throughout the Pleistocene compared to the relatively stable range of *N. obscura*, with the former spanning a range approximately twice as large as the latter during the mid-Pleistocene (Figure 4A). These patterns were similarly reflected within the standard deviations across timeslices per species, with *N. australis* showing much higher variation over a larger area (Figure 4C).

## Discussion

Our results demonstrate how spatial variation in demographic history may drive species-wide discordant responses to past climatic changes, even when local-scale impacts are concordant and species’ ecological traits are similar. Specifically, we show that within a shared climatic refugium for two co-distributed and ecologically similar freshwater fishes, demographic histories were largely concordant. However, towards the edges of this refugium demographic histories decreased in concordance, suggesting that range edge populations of *N. obscura* were more limited than *N. australis* in their capacity for expansion during more favourable climatic conditions. Together, our findings determine the importance of intraspecific, population-level dynamics in driving species-wide adaptation and resilience to climate change.

The temperate zone of southeast Australia has undergone significant environmental change since the Pliocene, owing to a combination of continent-wide aridification [24], eustatic changes [28] and major hydrological rearrangements [62]. These various aspects likely had significant impacts on the persistence and connectivity of freshwater lineages across the region [43]. This was supported by the high level of phylogenetic structure within *N. australis*, and the inferred migration pathways that correspond well to those previously suggested through ancient hydrological conduits [62]. Although phylogenetic patterns in *N. obscura* did not directly match the longitudinal gradient of populations, earlier phylogenetic analyses using allozymes and mitochondrial DNA showed a similar pattern [37]. This disjunction was attributed to potential historical connections from Mount Emu Creek into more western populations [35], although short branch lengths and low genetic diversity across the species may also indicate incomplete lineage sorting as a factor [63]. For both species, we denote two major clades: one of Murray-Darling Basin populations and another of coastal populations in *N. australis*, as suggested elsewhere [43], and two clades each containing two previously identified ESUs in *N. obscura* [37].

Within the species distribution models, a region of southwest Victoria was highlighted as a climatic refugium for both species throughout the Plio-Pleistocene. This region was consistently identified as suitable habitat for both species across all time slices. Although glacial maxima were associated with cold and arid conditions across Australia, coastal woodland habitats were likely buffered against intense aridification by oceanic circulation and relatively higher humidity and rainfall [64]. Other phylogeographical studies demonstrating limited impact of glacial maxima on connectivity supports the identity of this climatic refugium [27, 64]. Co-occurring populations within this shared refugium demonstrated highly congruent demographic histories at both more ancient (>1 Myr) and more recent (since the LGM) temporal scales. This concordance is expected when ecological traits, habitat preferences and environmental stability are shared across the species in question [21]. Although individual populations within each species demonstrated spatially variable demographic histories, comparisons across the two species showed similar patterns of *Ne* over time for most directly co-occurring populations.

Both pygmy perch species demonstrated temporally synchronous expansions during the LGM with post-glacial contractions across central populations. Despite intense inland aridification during glacial maxima, run-off in many southeast Australian rivers were likely much greater during the LGM [65]. These increased river flows have been attributed to seasonal snow melt of periglacial regions in the highlands and reduced vegetation cover, creating large rivers with enhanced run-off [65, 66]. Colder conditions and strong flows may have facilitated the observed concordant expansion in populations at this time, with the steep decline in flows during the early Holocene (14 – 7 Kya) potentially contributing to their more recent contraction [67]. However, concordance was reduced for pairwise populations that occurred closer to the edge of this shared refugium, suggesting the species had discordant responses at the fringe of the range. Similarly, phylogenetic patterns at the species-wide level varied between the two species, with clearer geographic sorting and historical migration across *N. australis* lineages compared to *N. obscura*.

Spatial variation in demographic history, and by extension concordance across taxa, may result from several different mechanisms [22, 68]. Particularly for narrowly distributed species, edge-of-range effects on populations close to the ecological tolerance threshold of the species may result in highly divergent patterns of demographic history and genetic diversity compared to more central populations [12, 68]. By extension, the ecological range of species may be a strong factor driving discordance when particular locations are at the periphery of the distribution of one species, but not another. Given the broad similarity in ecological traits between the two species and their co-occurring nature [69], it is unlikely that this discordance in species-wide responses to past climatic changes is a result of different ecologies.

However, some variation in microhabitat preference seems to exist between species, with *N. obscura* limited to larger, lowland channels and floodplains whereas *N. australis* is also found in streams and dense swamps [69]. This suggests greater habitat specialisation in *N. obscura*, which might drive lower SGV (or result from it) and impede range expansions. Thus, we cannot completely rule out some role of ecology and its interactions with genetic diversity in driving discordant responses. The lower genetic diversity in *N. obscura* could not be directly attributed to notable and widespread genetic bottlenecks, suggesting instead that the species suffered from a consistent pattern of being genetically depauperate. Combined, these factors suggest that long-term SGV may be a key factor driving the temporally and spatially widespread discordance in response to Pleistocene climate changes.

Adaptive responses, particularly in scenarios of range expansion, are often driven by soft sweeps of SGV [8]. While many studies focus on rapid adaptation from SGV in terms of invasive species colonising new habitats [70], similar dynamics can be expected to play a role in range expansions of native taxa [71]. In regard to range shifts across the Pleistocene, higher SGV may have predisposed *N. australis* to capitalise on the colder temperatures and stronger rivers of glacial periods and subsequently expand. Similarly, historical connectivity across now-isolated river drainages [28] likely facilitated interpopulation gene flow, which may have further bolstered SGV and adaptive potential [68, 72]. This gene flow in *N. australis* may have also facilitated range expansion if locally adaptive alleles were transferred into edge populations [70]. Contrastingly, a lack of long-term SGV within *N. obscura* may have prevented them from expanding under these conditions, leading to the species-wide discordance. The spatial variation in the degree of concordance, with discordance occurring at the edge of the *N. obscura* pre-glacial refugium, supports this conclusion.

Discordant species-wide responses to past climatic change may play an important role in contemporary genetic diversity and, by extension, current conservation efforts. For example, low genetic diversity resulting from historical bottlenecks can drive contemporary inbreeding depression [73]. Additionally, the parallels between historical range expansion scenarios and current reintroductions to conserve species demonstrates how historical processes may inform current practices [74]. For example, reduced adaptive capacity in *N. obscura* may have contributed to their local extirpation and to the failure of reintroductions of captive-born offspring at range margins, as documented for the lower Murray-Darling Basin [41]. This contrasts to the successful reintroduction of *N. australis* that simultaneously took place in that site using the same captive-breeding design [41, 42].

Understanding how, and which, species may be able to adapt under contemporary climate change remains a critical aspect of evolutionary biology [2]. Typically, this framework has focused on understanding how ecological traits may underpin individual species responses to climatic change [6]. However, demographic parameters are also critical components for species susceptibility to contemporary climate change [5]. Here, we demonstrate that intraspecific SGV may also be a critical component of species responses to climatic changes, particularly in range-edge populations. This corroborates studies indicating that adaptive potential is largely driven by SGV prior to the origination of major selective pressure [9] and suggests that considering broad ecology alone may not be enough to predict species’ ability to respond. Thus, understanding how the demographic history of individual populations may predispose, or hinder, species adaptive potential is an important component of conservation management of threatened species. For species with low SGV, proactive measures such as assisted gene flow and maintenance of effective population size may assist in their long-term conservation [75].

Long-term standing genetic variation drove discordance in the response of closely related and ecologically similar freshwater fishes to historical climate change, by facilitating range expansion of one species but not the other. However, in the centre of a shared habitat refugium, demographic histories were concordant, suggesting that spatial variation in the degree of concordance is linked to the interaction of standing genetic variation and distribution edge effects. Together, this demonstrates the importance of the maintenance of standing genetic variation for adaptive potential in response to climatic changes and the role of non-ecological traits in driving patterns of concordance or discordance.

## Acknowledgments

We acknowledge researchers that provided samples or participated in field expeditions, especially Mark Adams. This work was supported by an Australian Research Council grant (FT130101068 to L.B.B.) and by an Australian Government Research Training Program Scholarship to S.J.B. This work received logistic support from Flinders University, University of Canberra and the South Australian Museum.

