## Supplementary Material for "Variation in intraspecific demography drives localised concordance but species-wide discordance in responses to Plio-Pleistocene climatic change"

### Methods

*FastSimCoal2 individual population demographic modelling*

The demographic history of each population selected for comparative analysis (*n* = 3 per species) across the last glacial maximum (LGM) was estimated using five different models (Supplementary Figure S1). All populations were simulated using the same prior ranges for the same given model, with broad ranges to accommodate potential variation between populations. The best fitting model was determined using the best supported likelihood estimates to calculate the AIC:

$$\mathrm{AIC} = -2 \times \ln(model\ likelihood) + 2K,$$

where *K* refers to the number of parameters in the model, and Akaike weights:

$$\mathrm{AW} = e^{(-0.5 \times \Delta AIC)},$$

where ΔAIC refers to the difference in AIC for the given model compared to the lowest AIC model. The best supported model was then used to generate 100 independent SFS and re-estimate point estimates of parameters for the given model. The best parameter estimates per re-estimation were used to calculate the 95% confidence intervals of the inferred parameters. This bootstrapping process was automated by using a bash script (isaacovercast/fsc_2scripts/bootstrap_fsc.sh).

*Multi-DICE codemographic modelling*

We determined whether demographic histories were concordant across all six populations (3 co-occurring pairs) based on the aggregate site frequency spectrum (aSFS) within Multi-DICE [1]. The aSFS is an extension of other variants of the SFS which combines multiple SFS from separate populations, sorting them within each frequency bin in descending order across all taxa (and thus not preserving the original order of each SFS). Since the aSFS requires equal sample sizes across the combined taxa [2], individual SFS for all populations were down-projected to the smallest population sample size (*n* = 8 haploid samples) using the easySFS Python pipeline (github: isaacovercast/easySFS.git) and DaDi [3]. All populations were simulated under a fluctuating growth and bottleneck scenario following the results from individual FastSimCoal2 modelling. This included two epochs of exponential population size changes: an exponential growth followed by an exponential decline from this peak until the current day (looking forward in time). Broad priors for contemporary *Ne* (10K – 100K) and rate of population size changes ($\varepsilon 1$ = 5 – 20 and $\varepsilon 2$ = 0.1 – 1 for exponential decline and growth, respectively) were set to capture the array of results from individual FastSimCoal2 models. The timing of the change in population size patterns (from exponential growth to decline) was set ($\tau$) using a broad prior of 10 – 100 Kya to account for difficulties in detecting the timing of events. Population contractions were considered asynchronous if they occurred more than 1,000 years apart ($\beta$ = 1,000). We estimated co-demographic models in two separate stages, each consisting of a total of 1.5 million simulations. The simulated folded aSFS was estimated based on 1,000 SNPs, the approximate minimum number of variable sites used across the various individual population demography models in FastSimCoal2.

First, we allowed the proportion of synchronous taxa to vary within the model, ranging from one out of six ($\xi$ = 0.167) populations to all six ($\xi$ = 1) populations concordantly experiencing bottlenecks. The identifiability of this hyperparameter (i.e. the reliability with which it could be inferred) was determined using 50 pseudo-observed datasets and a confusion matrix, and the best supported hyperparameter value determined using the top 100 simulations in a Bayes Factor analysis. Second, we fixed ξ = 1 (based on the above results) and re-ran the model to better explore the remaining parameter values. Posterior distributions of important parameters (dispersion index of ε, mean ε across all populations, τ) were estimated using the top 100 simulations and the *abc* R package [4]. This approach is recommended by the developers of Multi-DICE [1].

*Species distribution modelling*

*Environmental data filtering*

All 19 bioclimatic variables from PaleoClim [5] and WorldClim [6] were cropped to the relevant extent. We filtered environmental variables based on a Pearson's correlation test of contemporary variables in SDMToolbox [7], removing any highly correlated variables ($r > |0.8|$). This filtering resulted in a final dataset of eight environmental layers - annual mean temperature (Bio1), mean diurnal range (Bio2), isothermality (Bio3), temperature seasonality (Bio6), mean temperature of the wettest quarter (Bio8), mean temperature of the driest quarter (Bio9), annual precipitation (Bio12) and precipitation seasonality (Bio15).

*Occurrence data filtering*

A total of 6,021 *N. australis* and 852 *N. obscura* occurrences were collated from various prior studies and the Atlas of Living Australia, and filtering to exclude coordinates with low spatial resolution or outliers that were beyond the bounds of the known distribution of either species. To reduce the impact of spatial autocorrelation, particularly in terms of potentially biased spatial sampling [8], we further reduced these datasets to a single occurrence per environmental raster cell. This resulted in the final datasets of 1,021 and 163 observations for *N. australis* and *N. obscura*, respectively

*Ensemble methods*

We generated three separate sets of pseudoabsences (*n* = 500) per species randomly from background cells >50km away from occurrences to reduce the likelihood of generating false absences within habitable areas [9]. Each dataset was replicated three times, with 80% of sites independently and randomly subset to train the model. SDMs were estimated for each dataset using the MaxEnt model, Random Forest (RF) and a generalised linear model (GLM; *n* = 27 models total). Each model was evaluated using both the area under the receiver operating curve (ROC) and the true skill statistic (TSS). Ensemble models per time period and species were generated using the weighted mean of all models with a TSS > 0.7. Despite this threshold, no models were excluded from the ensemble.

For quantitative assessment of distribution dynamics over time, we estimated the mean and standard deviation of suitability per cell across all ten time period ensembles per species. To assess changes in area over time, we first converted each ensemble SDM – as well as each of the 27 individual models – to binary presence-absence maps by first calculating a minimum presence threshold based on the minimum suitability of the top 90% of putative occurrences under current conditions (derived from https://babichmorrowc.github.io/post/2019-04-12-sdm-threshold/, accessed 10/10/2019). Each of these thresholds was then applied across all projections for the given method (*n* = 28 thresholds total). The area of each binary raster was estimated by counting the number of inferred presence cells: given that all rasters were of the same extent and resolution, these values were directly comparable across all approaches.

**Table S1:** Locality data for samples used in this study. Abbreviations described in the table were those used for further analyses, while *n* refers to the number of individuals sequenced per population. *Nannoperca vittata* samples were only included as an outgroup in the phylogenetic analysis.

| Species | Population | Abbreviation | Field code | Latitude | Longitude | N |
|---|---|---|---|---|---|---|
| *N. australis* | Angas, Middle Ck. Junction | NauANG | F-FISH84 | -35.25 | 138.887 | 5 |
| | Lake Alexandrina | NauALE | SPPBrA* | -35.395 | 139.008 | 4 |
| | Glenelg Rvr., Glenisla | NauGRG | F-FISH78: PU0014-SPP | -37.1369 | 142.262 | 5 |
| | Meriri Rvr., Grassmere | NauMRG | F-FISH78: PU00-22SPP | -38.2638 | 142.5151 | 5 |
| | Curdies Rvr., Curdie | NauCRC | F-FISH78: PU00-24SPP | -38.5189 | 142.836 | 4 |
| | Gellibrand Rvr. floodplain | NauGRF | F-FISH97: PU02-92SPP | -38.6919 | 143.1675 | 5 |
| | Barongarook Ck., Colac | NauBAR | SPP08-13 | -38.36 | 143.6404 | 4 |
| | Mundy Gully | NauMG | F-FISHY8: PU08-11SPP | -37.9523 | 143.3794 | 4 |
| | Gnarkeet Ck., Hamilton | NauGCH | F-FISHY2: PU00-27SPP | -37.9706 | 143.4674 | 4 |
| *N. obscura* | Cleland, Hamilton Island | NobCHI | YPPBr* | -35.537 | 138.905 | 5 |
| | Mt. Emu Ck. | NobMEC | PU02-112YPP | -38.325 | 142.759 | 5 |
| | Mosquito Ck., Langkoop | NobMCL | PU00-16YPP | -37.0938 | 140.9841 | 4 |
| | Shaw Rvr. | NobSR | PU02-113YPP | -38.1699 | 142.0902 | 2 |
| | Merri Rvr., Grassmere | NobMRG | PU02-111YPP | -38.275 | 142.542 | 4 |
| | Gnarkeet Ck., Lismore | NobGCL | PU00-27YPP | -37.9102 | 143.4469 | 5 |
| | Deep Ck., Lancefield | NobDee | F-FISH97: PU02-106YPP | -37.259 | 144.713 | 2 |
| *N. vittata* | Chesapeake Rvr. | *Outgroup* | F-FISHx2: V22, V64, V83, V85, V185 | -34.8362 | 116.333 | 5 |
| **Total** | | **16** | | | | **72 (77)** |

**Table S2**: Initial prior ranges for individual population demographic models estimated within FastSimCoal2. All priors were set using uniform distributions, with the same conditions across all six populations for the same model. The relationship of parameters to the model, and broad overviews of the models themselves, are demonstrated in Supplementary Figure S1.

| **Parameter** | **Model** | | | | |
|---|---|---|---|---|---|
| | **1** | **2** | **3** | **4** | **5** |
| NCURR | $10^3$ - $10^4$ | $10^3$ - $10^4$ | $10^3$ - $10^4$ | $10^3$ - $10^4$ | $10^3$ - $10^4$ |
| RATE | $-10^{-4}$ – 0 | 0 – $10^{-4}$ | NA | $-10^{-4}$ – 0 | 0 – $10^{-4}$ |
| T1 | 18,000 – 21,000 | 18,000 – 21,000 | NA | 18,000 – 21,000 | 18,000 – 21,000 |
| R2 | NA | NA | NA | 0 – $10^{-4}$ | $-10^{-4}$ – 0 |
| D2 | NA | NA | NA | 1,000 – 20,000 | 1,000 – 20,000 |
| T2 | NA | NA | NA | = T1 + D2 | = T1 + D2 |

**Table S3:** Pearson's pairwise correlation for all 19 bioclimatic variables obtained from WorldClim v1.4. Uncorrelated variables (|R| ≤ 0.8) are highlighted in bold, and were subsequently used in species distribution modelling.

| **Variable** | **Bio1** | **Bio2** | **Bio3** | Bio4 | Bio5 | **Bio6** | Bio7 | **Bio8** | **Bio9** | Bio10 | Bio11 | **Bio12** | Bio13 | Bio14 | **Bio15** | Bio16 | Bio17 | Bio18 | Bio19 |
|---|---|---|---|---|---|---|---|---|---|---|---|---|---|---|---|---|---|---|---|
| **Bio1** | | | | | | | | | | | | | | | | | | | |
| **Bio2** | **0.72** | | | | | | | | | | | | | | | | | | |
| **Bio3** | **0.08** | **0.06** | | | | | | | | | | | | | | | | | |
| Bio4 | 0.60 | 0.83 | -0.47 | | | | | | | | | | | | | | | | |
| Bio5 | 0.93 | 0.89 | -0.05 | 0.82 | | | | | | | | | | | | | | | |
| **Bio6** | **0.48** | **-0.16** | **0.41** | -0.36 | 0.17 | | | | | | | | | | | | | | |
| Bio7 | 0.69 | 0.94 | -0.24 | 0.96 | 0.89 | -0.28 | | | | | | | | | | | | | |
| **Bio8** | **0.62** | **0.46** | **0.04** | 0.40 | 0.59 | **0.30** | 0.44 | | | | | | | | | | | | |
| **Bio9** | **0.51** | **0.35** | **0.10** | 0.23 | 0.46 | **0.30** | 0.31 | -0.19 | | | | | | | | | | | |
| Bio10 | 0.95 | 0.83 | -0.10 | 0.79 | 0.98 | 0.25 | 0.84 | 0.60 | 0.47 | | | | | | | | | | |
| Bio11 | 0.88 | 0.43 | 0.37 | 0.18 | 0.69 | 0.79 | 0.30 | 0.52 | 0.52 | 0.73 | | | | | | | | | |
| **Bio12** | **-0.80** | **-0.75** | **-0.15** | -0.57 | -0.82 | **-0.24** | -0.69 | **-0.51** | **-0.42** | -0.80 | -0.66 | | | | | | | | |
| Bio13 | -0.78 | -0.77 | -0.10 | -0.61 | -0.82 | -0.18 | -0.72 | -0.51 | -0.41 | -0.80 | -0.61 | 0.98 | | | | | | | |
| Bio14 | -0.76 | -0.63 | -0.31 | -0.37 | -0.71 | -0.39 | -0.51 | -0.38 | -0.50 | -0.70 | -0.73 | 0.91 | 0.84 | | | | | | |
| **Bio15** | **-0.22** | **-0.35** | **0.36** | -0.54 | -0.35 | **0.27** | -0.47 | **-0.36** | **0.09** | -0.34 | 0.05 | **0.21** | 0.35 | -0.14 | | | | | |
| Bio16 | -0.79 | -0.77 | -0.09 | -0.63 | -0.83 | -0.18 | -0.73 | -0.54 | -0.38 | -0.81 | -0.61 | 0.98 | 1.00 | 0.84 | 0.36 | | | | |
| Bio17 | -0.77 | -0.67 | -0.26 | -0.43 | -0.74 | -0.34 | -0.57 | -0.40 | -0.49 | -0.73 | -0.71 | 0.94 | 0.88 | 0.99 | -0.09 | 0.87 | | | |
| Bio18 | -0.71 | -0.60 | -0.28 | -0.35 | -0.67 | -0.35 | -0.49 | -0.23 | -0.62 | -0.66 | -0.69 | 0.87 | 0.81 | 0.95 | -0.14 | 0.79 | 0.95 | | |
| Bio19 | -0.75 | -0.76 | -0.07 | -0.62 | -0.81 | -0.15 | -0.72 | -0.61 | -0.27 | -0.78 | -0.57 | 0.96 | 0.98 | 0.79 | 0.41 | 0.99 | 0.83 | 0.71 | |

**Table S4**: Population genetic summaries of *N. australis* and *N. obscura.* Populations with $n \leq 3$ were excluded due to low sample size. Where applicable, values are reported as means of means across all loci ± standard deviation under rarefaction ($n = 4$ samples) using the species-wide alignment SNPs. For Lake Alexandrina populations (NauALE and NobCHI), gene diversities were also calculated for individual population alignments and rarefaction of $n = 15$ samples to take advantage of larger sample sizes (reported second). Totals are reported as species-level averages across the alignment(s), excluding SNPs which refers to the total number of SNPs in the species-wide alignment.

| **Alignment** | | | **Species-wide** | | **Individual populations** | | |
|---|---|---|---|---|---|---|---|
| **Species** | **Population** | **N** | **AR** | **Hs** | **Ne** | **Π** | **SNPs** |
| ***N. australis*** | NauANG | 5 | 1.138 (± 0.339) | 0.029 (± 0.106) | | $1.59 \times 10^{-4}$ | 2,446 |
| | NauALE | 5<br>20 | 1.354 (± 0.480) | 0.062 (± 0.165)<br>0.179 (± 0.142) | 71.3 [35.9 – 520.9] | $1.23 \times 10^{-3}$ | 1,198 |
| | NauGRG | 5 | 1.123 (± 0.313) | 0.035 (± 0.108) | | $8.84 \times 10^{-5}$ | 5,282 |
| | NauMRG | 5 | 1.116 (± 0.312) | 0.037 (± 0.119) | | $1.62 \times 10^{-4}$ | 3,835 |
| | NauGRF | 5 | 1.092 (± 0.285) | 0.027 (± 0.106) | | $7.84 \times 10^{-5}$ | 2,969 |
| | NauBAR | 4 | 1.240 (± 0.428) | 0.061 (± 0.145) | | $2.02 \times 10^{-4}$ | 5,084 |
| | NauMG | 4 | 1.216 (± 0.413) | 0.056 (± 0.138) | | $1.57 \times 10^{-4}$ | 5,357 |
| | NauGCH | 4 | 1.181 (± 0.387) | 0.047 (± 0.136) | | $1.88 \times 10^{-4}$ | 4,272 |
| | **Total** | 56 | 1.241 | 0.0439 | | $2.83 \times 10^{-4}$ | 17,389 |
| ***N. obscura*** | NobCHI | 5<br>15 | 1.151 (± 0.347) | 0.038 (± 0.117)<br>0.136 (± 0.108) | 14.9 [4.6 – 652.2] | $1.69 \times 10^{-4}$ | 721 |
| | NobMEC | 5 | 1.030 (± 0.166 | 0.011 (± 0.069) | | $2.11 \times 10^{-5}$ | 1,002 |
| | NobMCL | 4 | 1.100 (± 0.290) | 0.026 (± 0.094) | | $4.18 \times 10^{-5}$ | 1,633 |
| | NobMRG | 4 | 1.058 (± 0.234) | 0.020 (± 0.093) | | $3.23 \times 10^{-5}$ | 1,454 |
| | NobGCL | 5 | 1.070 (± 0.250) | 0.019 (± 0.088) | | $2.49 \times 10^{-5}$ | 1,350 |
| | **Total** | 33 | 1.168 | 0.0237 | | $5.78 \times 10^{-5}$ | 15,715 |
| **T-test** | | | T = 2.38<br>($p < 0.04$) | T = 2.93<br>($p = 0.01$) | | W = 56<br>($p = 0.01$) | T = 4.63<br>($p < 0.01$) |

**N** = total number of samples per population. **AR** = rarefied allelic richness. **Hs** = rarefied gene diversity. ***Ne*** = effective population size, with 95% confidence intervals estimated by jack-knifing in square brackets. **π** = nucleotide diversity. Populations means across species were compared using either an unpaired two-samples t-test (**T**) or Wilcoxon rank test (**W**).

**Table S5:** Posterior distributions of parameters from co-demographic models in Multi-DICE. Posterior probabilities were calculated using the top 0.0067% (*n* = 100) simulations out of 1.5M total simulations. **τ** = timing of population size change (bottleneck). **ε1** = magnitude of exponential population decline. ***Ne*** = current effective population size. **Mean(ε)** = average of ε across all six populations. **Ω(ε)** = dispersion index of ε (Var(ε)/Mean(ε)). Taxon-specific parameters (ε1 and Ne) are reported as the min – max range across all six populations.

| | Parameter | | | | |
|---|---|---|---|---|---|
| | τ | ε1 | *Ne* | Mean(ε1) | Ω(ε1) |
| Min | 73,751 | 5.009 – 5.061 | 10,026 – 10,542 | 5.673 | 0.019 |
| 2.5% | 77,073.5 | 5.083 – 5.218 | 10,155 – 11,554 | 6.393 | 0.152 |
| Mean | 91,452.34 | 8.438 – 8.734 | 22,212 – 24,887 | 8.618 | 1.41 |
| 97.5% | 99,851.5 | 16.012 – 18.843 | 58,988 – 72,268 | 11.390 | 2.991 |
| Max | 99,971 | 17.606 – 19.996 | 82,156 – 99,863 | 12.575 | 3.338 |

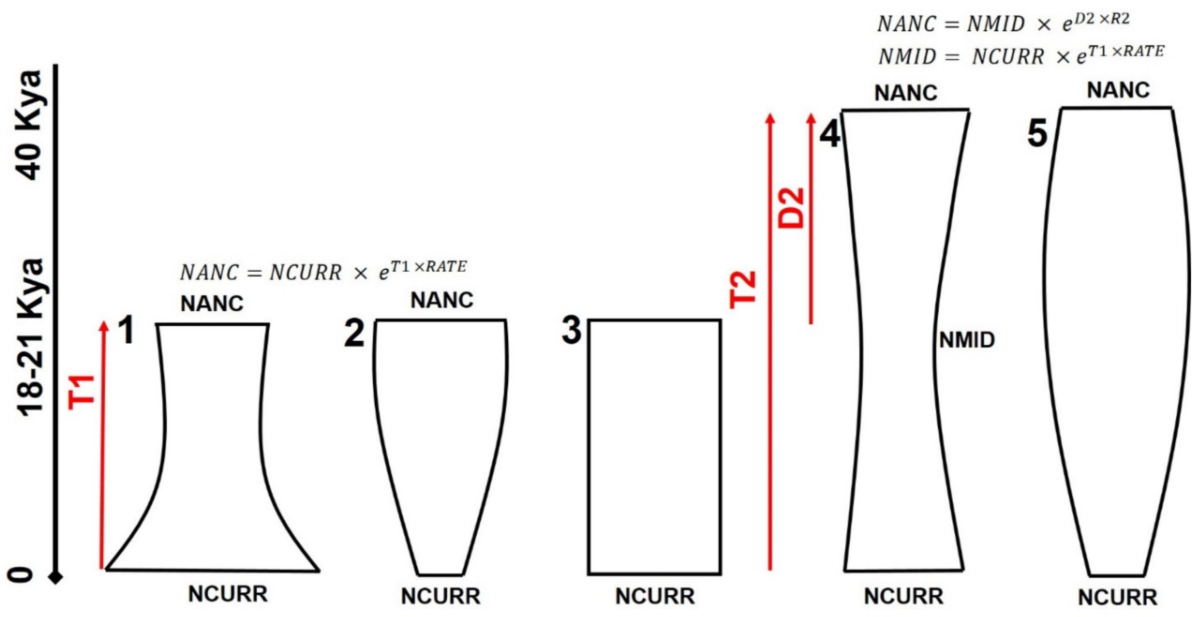


**Figure S1:** Diagrammatic representations of demographic syndromes tested per population selected for codemographic analysis, using FastSimCoal2. Each corresponds to the same numbers in Results and other Supplementary Figures. The width of each diagram corresponds to *Ne* over time, changing from current *Ne* (at the bottom of the figure) back in time (moving upwards). *RATE* and *R2* parameters indicate rates used to calculate exponential growth or decay of *Ne* over time, with formula for *Ne* at given points indicated above each model. Red parameters indicate timing of the end of growth or decay rates, depending on the model.

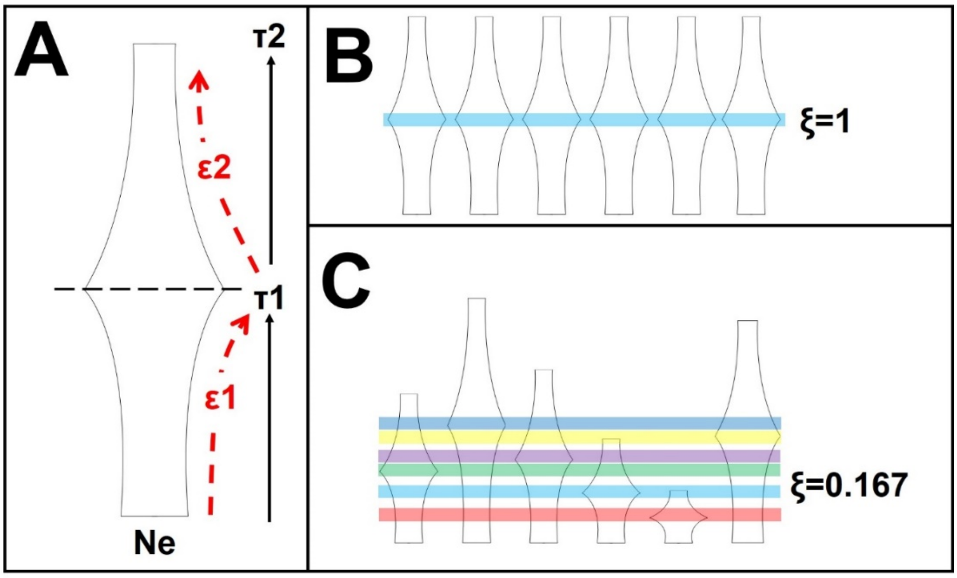


**Figure S2:** Diagrammatic representations of codemographic models within Multi-DICE. **A**) Example of parameters within a single taxon of the demographic model, containing two population size changes over time. A single broad prior distribution is set for parameters across all taxa, but posteriors may vary freely across taxa within the model. Arrows for τ parameters indicate direction backwards in time (i.e. τ1 is more recent in the past compared to τ2). Parameter priors are described in the Methods section of the Supplementary Material. **B)** An example of a fully synchronous model, when the hyperparameter ξ (the proportion of taxa belonging to a single synchronous event) is one. All populations share the timing of the population size change event, within the range of buffer β. **C**) An example of a fully asynchronous model, when ξ = 1/6 taxa per event. All populations have different timings of events, with each event >β generations from one another. The hyperparameter allows intermediate variations of ξ, which are not demonstrated here. We used a set of models to first determine the most likely value of ξ, before fixing this hyperparameter to better explore the remaining parameters. All other parameters and priors were kept constant between the two steps.

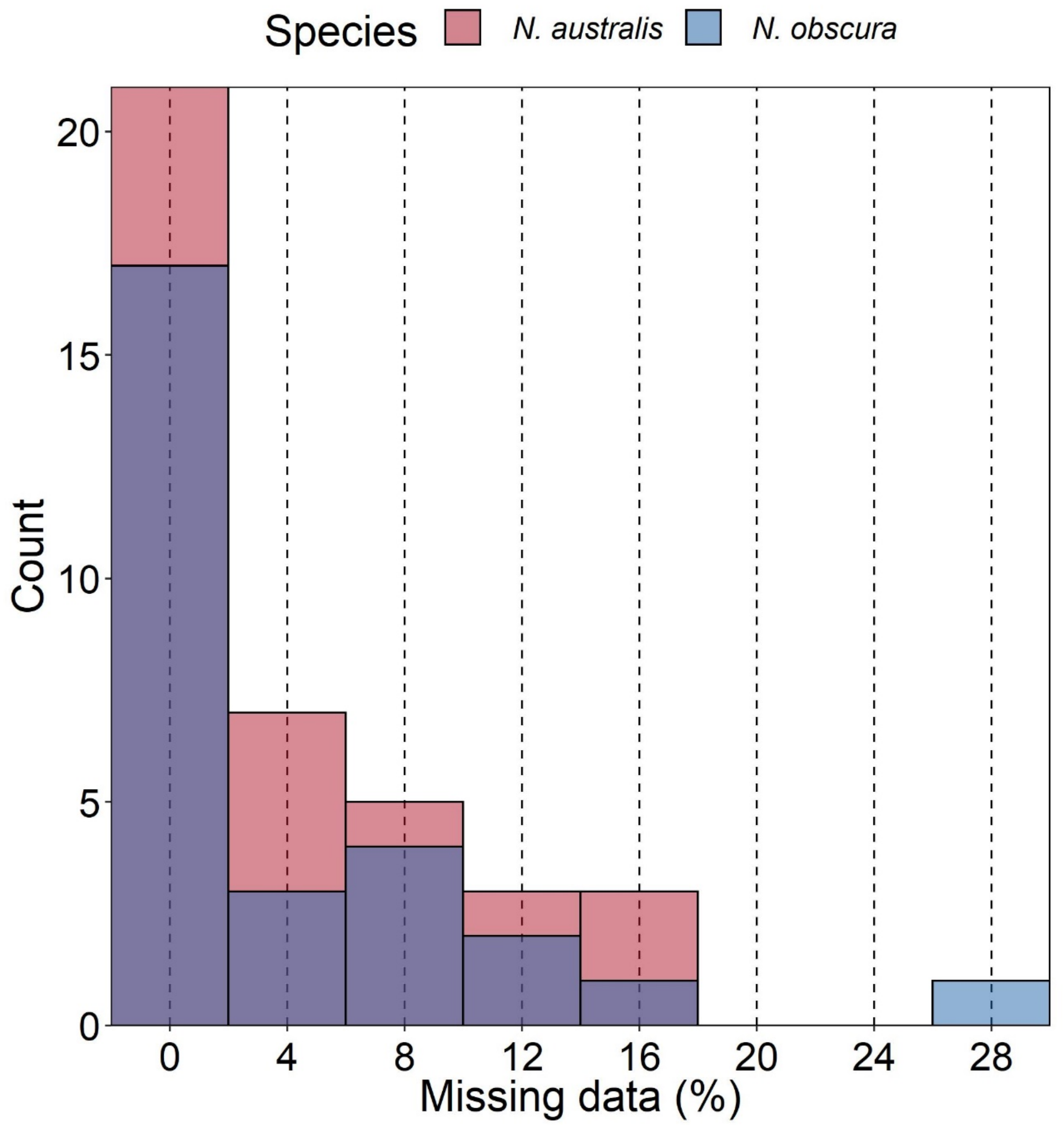


**Figure S3:** Histogram of missing data per sample in the species-wide alignments. Bar colours denote species.

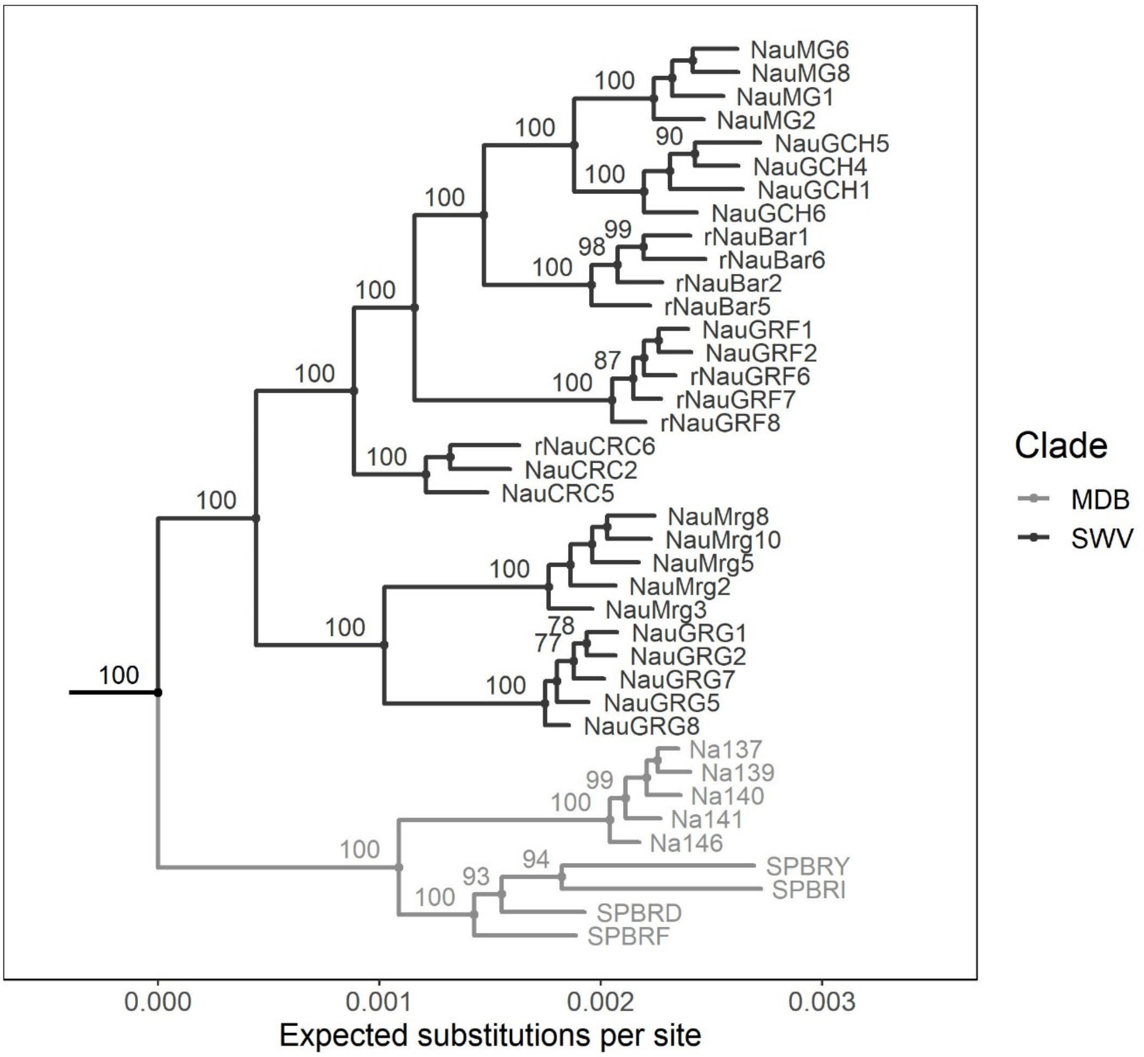


**Figure S4:** Full maximum likelihood phylogenetic tree for *N. australis* estimated using RAxML and based on 19,428 concatenated ddRAD loci. The tree was rooted using *N. vittata* as the outgroup. Node values show bootstrap support under 1,000 RELL bootstraps (only nodes with >75% support reported). Branch colours indicate the basin of origin for each clade (MDB = Murray-Darling Basin, SWV = southwest Victoria).

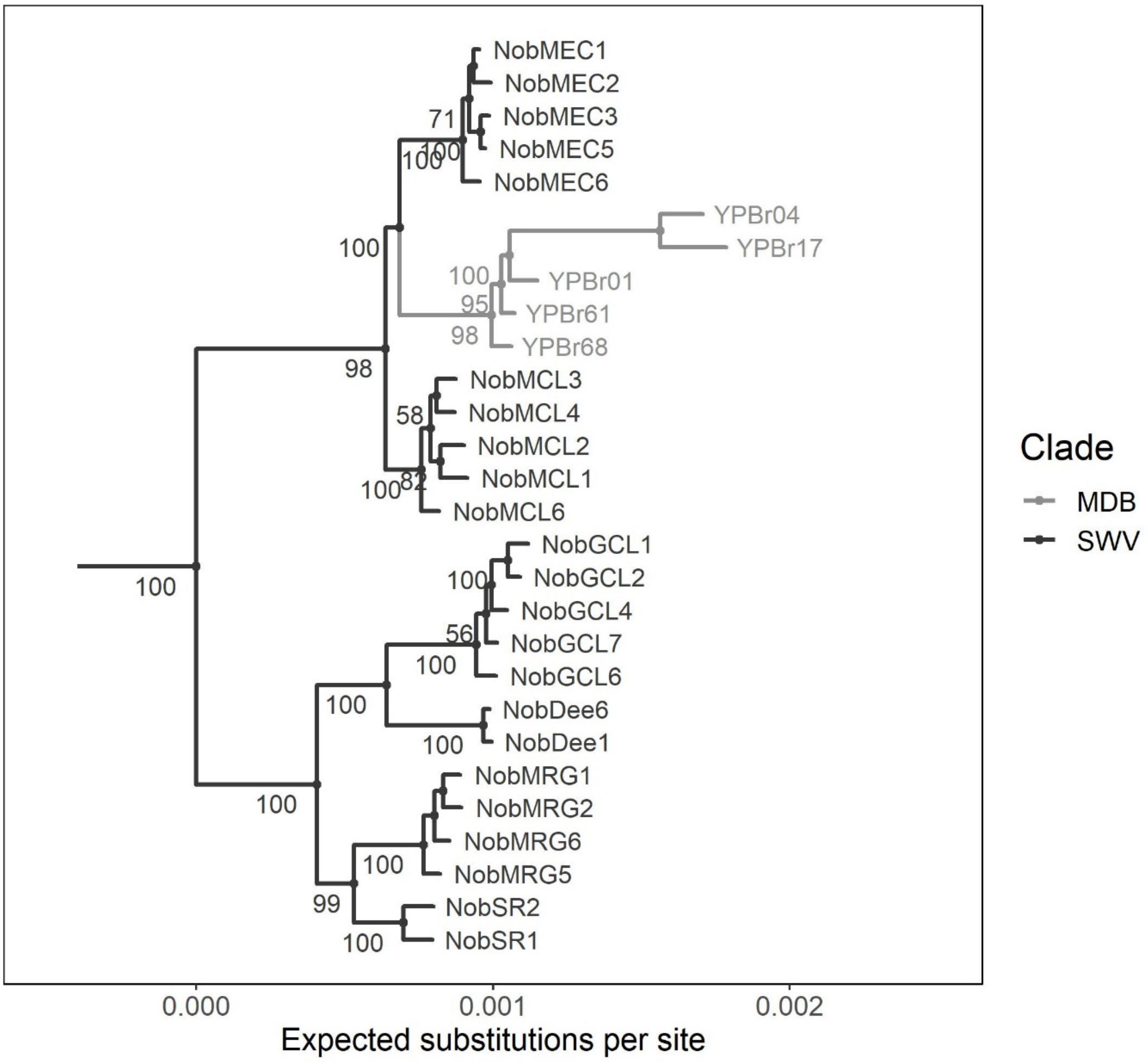


**Figure S5**: Full maximum likelihood phylogenetic tree for *N. obscura* estimated using RAxML and based on 12,705 concatenated ddRAD loci. The tree was rooted using *N. vittata* as the outgroup. Node values show bootstrap support under 1,000 RELL bootstraps (only nodes with >50% support reported). Branch colours indicate the basin of origin for each clade (MDB = Murray-Darling Basin, SWV = southwest Victoria).

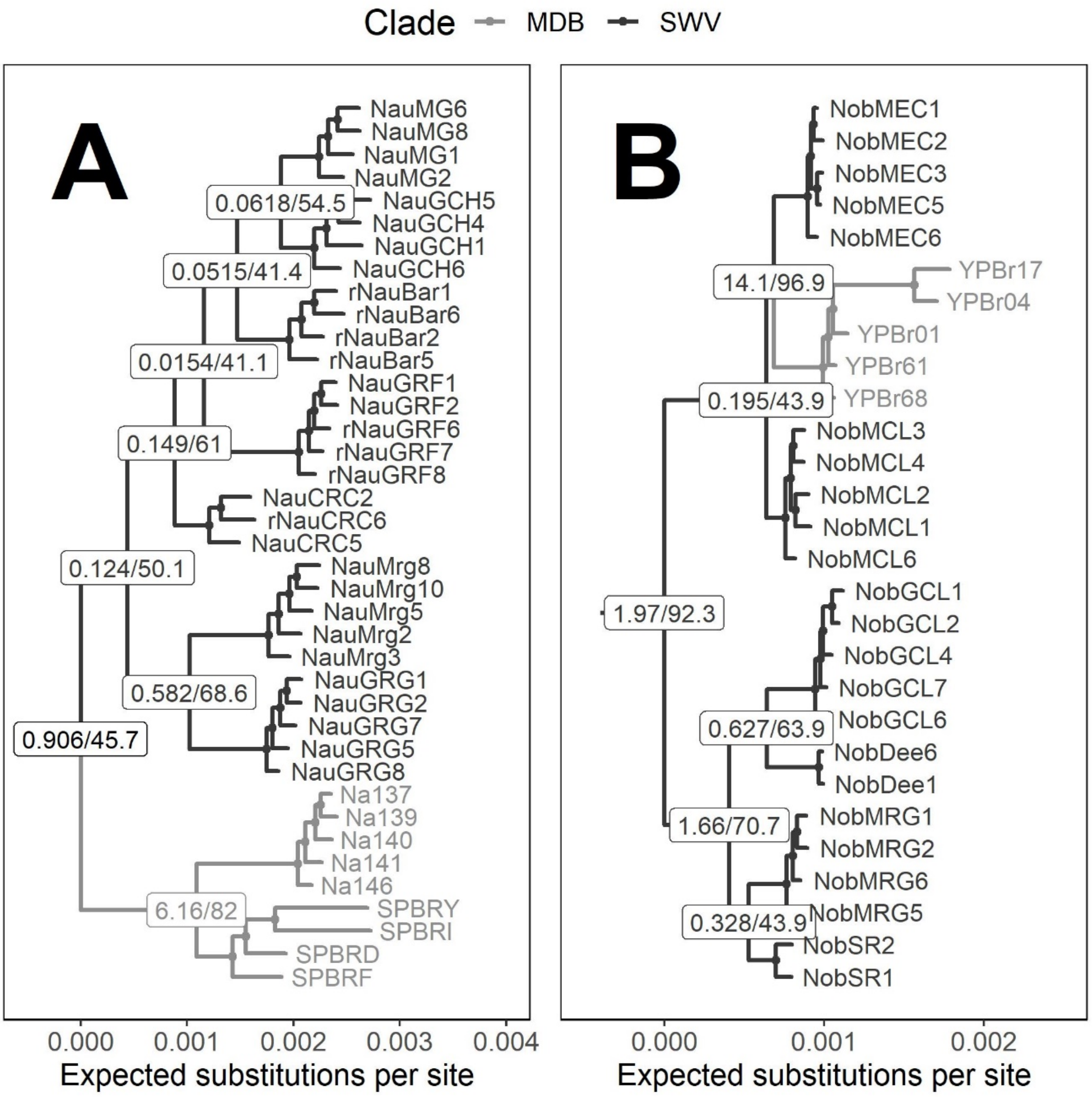


**Figure S6:** Gene and site concordance factors between gene trees estimated with individual RAD loci and the concatenated phylogeny for *N. australis* (**A**) and *N. obscura* (**B**). Population-level divergences and above are labelled with concordance factors, with gene concordance factors reported first and site concordance factors reported second. Both factors are scaled out of 100.

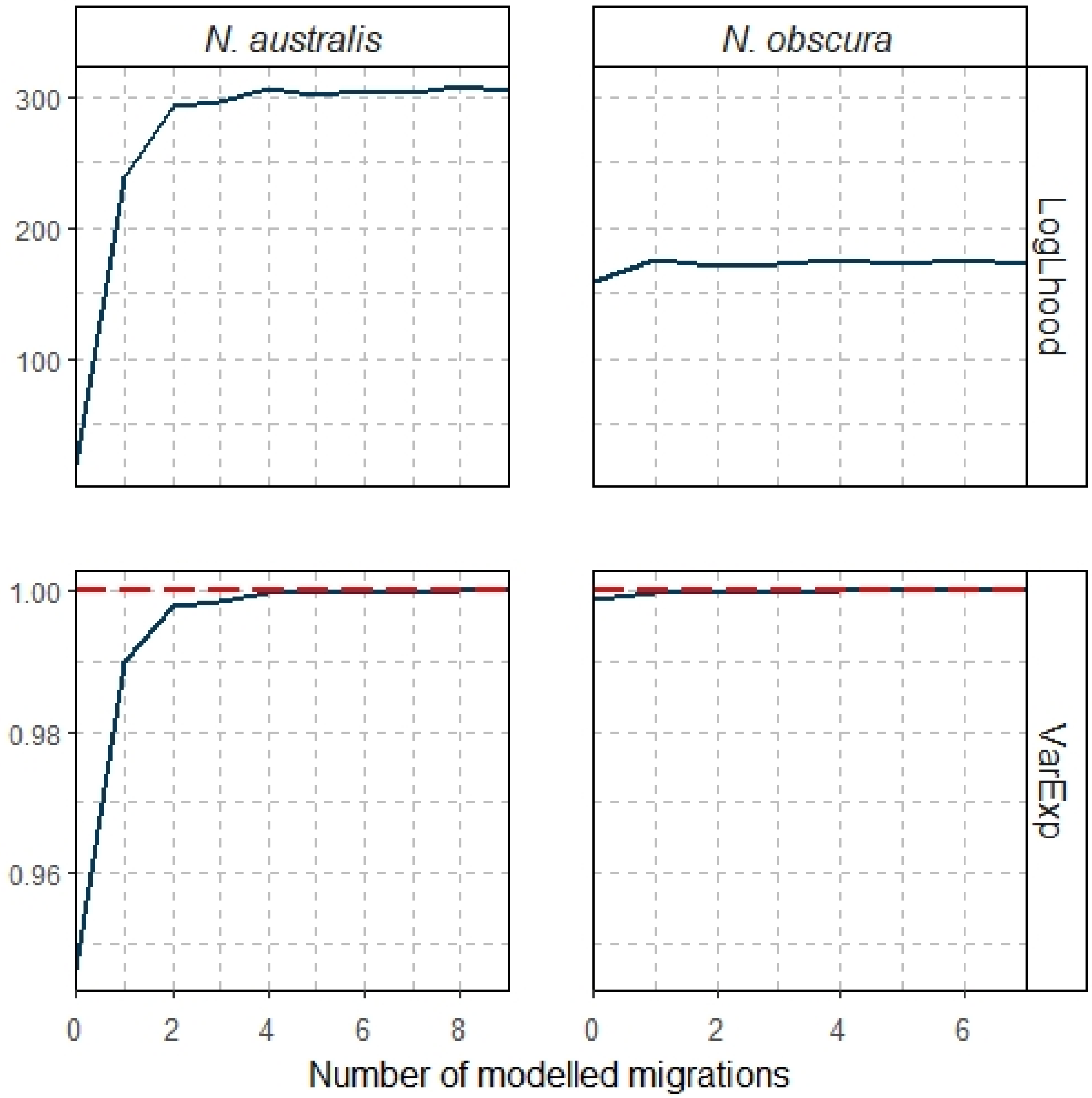


**Figure S7:** Log likelihoods (top) and percentage of variation explained (bottom) of population mixtures and splits modelled with varying numbers of migration edges in TreeMix. Dashed red line indicates the asymptote of the variation explained.

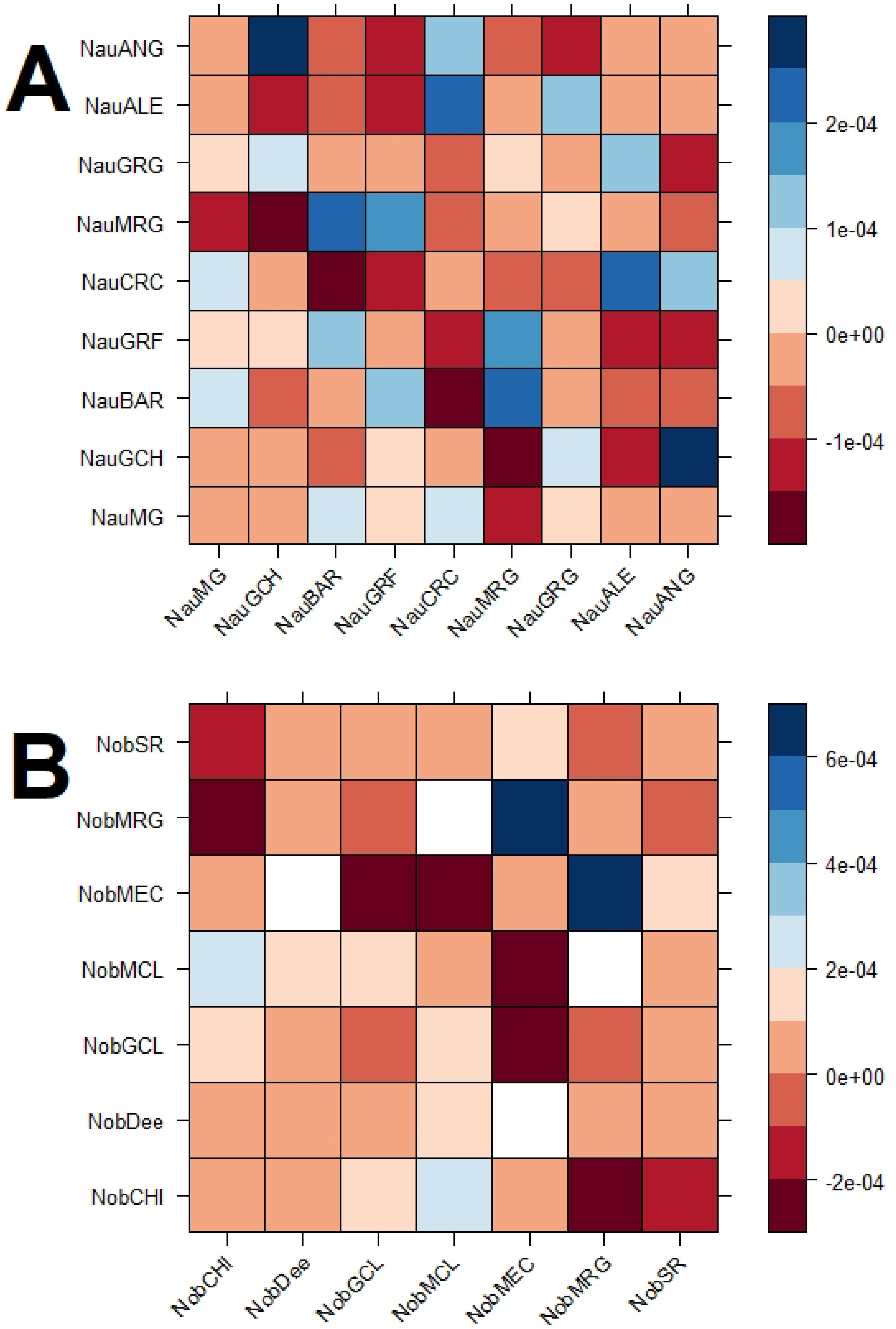


**Figure S8:** Residual matrices for **A)** *N. australis* and **B)** *N. obscura* under the best supported TreeMix models.

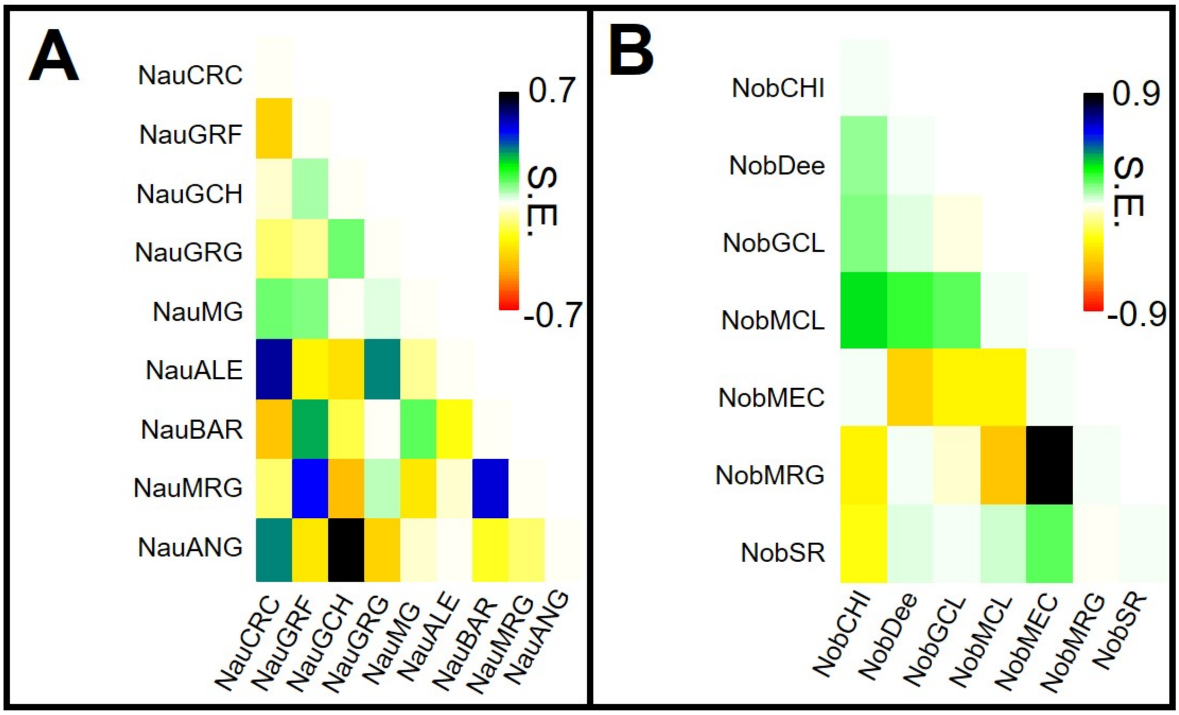


**Figure S9**: Standard errors of the covariance matrices for **A)** *N. australis* and **B)** *N. obscura* under the best supported TreeMix models.

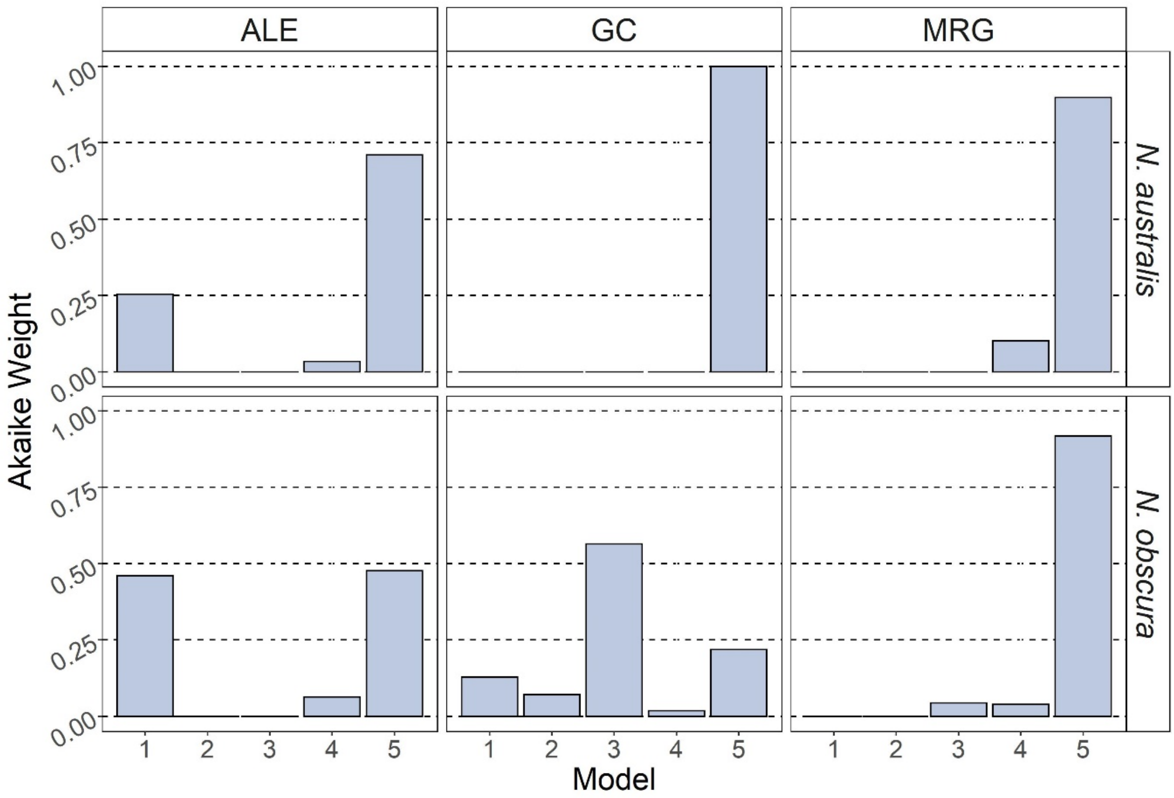


**Figure S10:** Akaike weights for demographic syndrome models using FastSimCoal2. Individual demographic models relate to the descriptions and numbering in Figure S1.

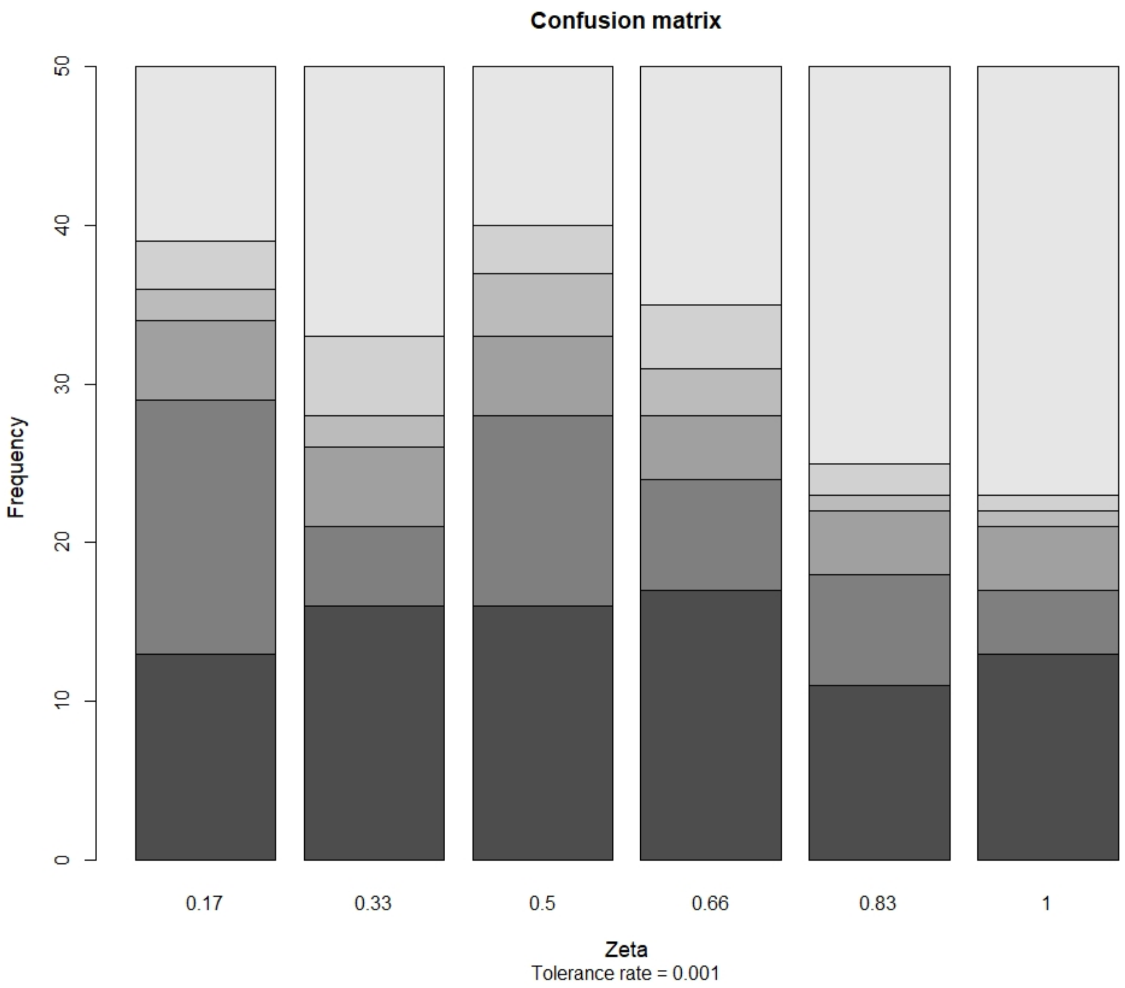


**Figure S11:** Confusion matrix of ξ hyperparameter within the co-demographic Multi-DICE model, estimated using 50 pseudo-observed datasets and the top 1,500 simulations (out of 1.5M simulations total). Colours range from dark grey (ξ = 0.17) to light grey (ξ = 1) for each possible value of ξ.

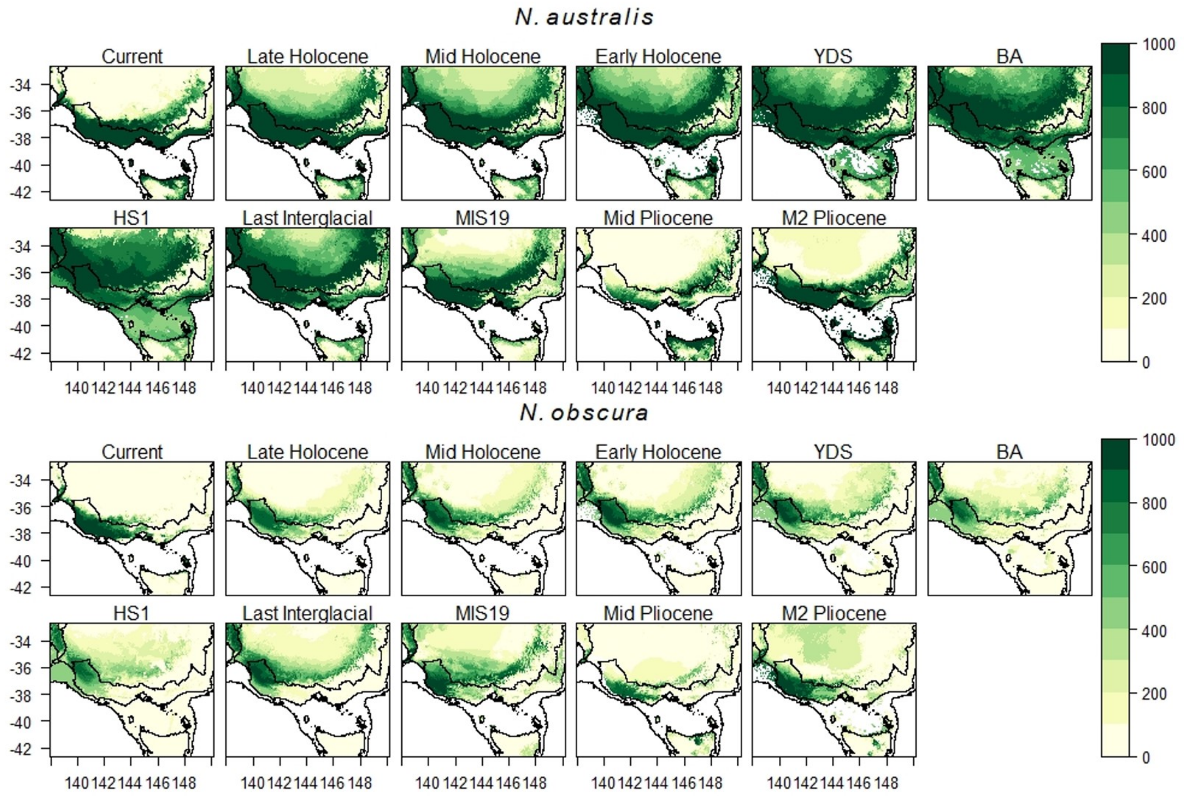


**Figure S12:** Ensemble species distribution models projected from current conditions to the mid-Pliocene for both *N. australis* and *N. obscura*. SDMs are based on 1,021 and 163 occurrences, respectively, and 10 environmental variables (9 bioclimatic variables + elevation).
